# Nitrogen dynamics and fixation control cyanobacterial abundance, diversity, and toxicity in Lake of the Woods (USA, Canada)

**DOI:** 10.1101/2023.04.12.536566

**Authors:** Kaela E. Natwora, Adam J. Heathcote, Mark B. Edlund, Shane E. Bowe, Jake D. Callaghan, Cody S. Sheik

## Abstract

Our understanding of drivers of cyanobacterial harmful algal blooms (cHABs) is evolving, but it is apparent that not all lakes are created equal. Nitrogen (N) is an important component of all cHABs and is crucial for cyanotoxin production. It is generally assumed that external nitrogen inputs are the primary N source for cHABs. However, in northern lakes, nitrogen inputs are typically low, and suggests that internal nitrogen cycling, through heterotrophic organic matter decomposition or nitrogen fixation, may play a significant role in cHAB development and sustainment. Using Lake of the Woods as a testbed, we quantified nutrients, cyanotoxins, nitrogen fixation, and the microbial community in the southern extent of the lake. During our temporal study, inorganic nitrogen species (NO_3_^-^+NO_2_^-^ and NH_4_^+^) were either at very low concentrations or below detection, while phosphorus was in excess. These conditions resulted in nitrogen-deficient growth and thereby favored nitrogen fixing cyanobacterial species. In response, nitrogen fixation rates increased exponentially throughout the summer and coincided with the *Aphanizomenon* sp. bloom. Despite nitrogen limitation, microcystin, anatoxin, saxitoxin, and cylindrospermopsin were all detected, with microcystin being the most abundant cyanotoxin detected. Microcystin concentrations were highest when free nitrogen was available and coincided with an increase in *Microcystis.* Together, our work suggests that internal nitrogen dynamics are responsible for the dominance of nitrogen fixing cyanobacteria and that additions of nitrogen may increase the likelihood of other cyanobacterial species, currently at low abundance, to increase growth and cyanotoxin production.

**Statement of Significance:** This study is the first assessment of nitrogen fixation rates and water column 16S rRNA gene amplicon sequencing in Lake of the Woods during a harmful algal bloom season. The aim of this study is to better understand nitrogen dynamics and the microbial ecology of cyanobacterial harmful algal blooms on Lake of the Woods. Result from this study reveal that internal nitrogen cycling via nitrogen fixation may alleviate nitrogen deficiencies, and structure and control the cyanobacterial community and cyanotoxin production. Molecular analysis reveals that cyanotoxins in Lake of the Woods are produced by less abundant cyanobacteria that are limited by nitrogen. This study has significant management implication as agencies continue to mitigate toxic blooms on Lake of the Woods, the largest shoreline lake in the United States. Our work is an important initial assessment and jumping off point for further research on Lake of the Woods when assessing how nitrogen plays a role in bloom formation and toxicity. Submitting to L&O, we believe would allow for the greatest outreach and access to an audience that will continue to build upon our findings. Additionally, submitting with L&O our work will reach beyond the scientific audience, but also reach other parties participating in the mitigation of harmful algal blooms.

## Introduction

Cyanobacterial harmful algal blooms (cHABs) are increasing in frequency and duration worldwide and pose a continued threat to all freshwater ecosystems (D. M. Anderson, 2009; Gobler, 2020; G. Hallegraeff *et al*., 2003; Ho *et al*., 2019). Many species of cyanobacteria are capable of producing potent suites of toxins (Carmichael, 1997) that have far-reaching effects on ecosystem services (Bauer *et al*., 2010; G. M. Hallegraeff *et al*., 2004; Landsberg, 2002; Young *et al*., 2020). Aquatic life, such as waterfowl or mammals that depend upon these affected waters, has seen mass mortalities and diseases resulting from cyanotoxin consumption and exposure (Landsberg, 2002). Likewise, domestic animal mortality has also been associated with rising cHABs (Hilborn & Beasley, 2015). Recent cHABs in Lake Erie showcases the devastating and costly effects that blooms can have on cities that depend on drinking water reservoirs prone to toxin producing blooms. The Toledo Drinking Water Crisis, caused by microcystin-producing *Microcystis spp.* (Steffen *et al*., 2017), left more than 400,000 citizens without drinking water and cost millions of dollars to remediate (Jetoo *et al*., 2015). While *Microcystis* spp. are abundant and common sources of microcystin, there are many cyanobacterial genera capable of synthesizing toxins (i.e., microcystin, anatoxin, saxitoxin, and cylindrospermopsin), including *Dolichospermum*, *Aphanizomenon, Planktothrix* and *Cylindrospermopsis* (Codd *et al*., 2005). While, we generally understand that nutrient availability plays a large role in driving blooms, several questions remain as to what combination of biotic and abiotic factors initiate and prolong toxin production during the course of a cHAB (Pal *et al*., 2020).

Nutrient availability, primarily phosphorus (P), nitrogen (N), or a combination of the two, are common drivers of cyanobacterial bloom initiation and toxin production (Paerl *et al*., 2016; Schindler, 1974). However, the phosphorous only nutrient paradigm does not adhere to all lakes, as we have seen blooms in oligotrophic systems with highly imbalanced N:P ratios become more prevalent (Reinl *et al*., 2021; Sheik *et al*., 2022; Sterner *et al*., 2020). Nonetheless, biological preference for various forms of N (oxidized *vs* reduced) is species dependent, and the bioavailability of P is likely dependent on the mineral form and solubility, which are pH dependent. Furthermore, N:P ratios of aquatic systems are a continuum over a growing season (H. Li *et al*., 2020; Y. Li *et al*., 2021) and may display periods of nitrogen- or phosphorus-deficient growth (Guildford & Hecky, 2000). Thus, the seasonal development of blooms, i.e., their phenology, is dependent on the consortia of organisms present, their metabolic needs, and in-lake nutrient dynamics.

While phosphorus is important and necessary for growth, molar nitrogen concentrations are always greater than phosphorus within a cell (Guildford & Hecky, 2000; Redfield, 1958), and nitrogen availability regulates primary production in both marine and freshwater environments (Conley *et al*., 2009; Howarth & Marino, 2006; Paerl *et al*., 2016). Often, inorganic and organic nitrogenous compounds are introduced to riverine and lake systems through watershed runoff, atmospheric deposition, groundwater, or sediment resuspension (Gardner *et al*., 2006; Lepori & Keck, 2012; Vitousek *et al*., 1997). To date, efforts to mitigate blooms have focused on limiting inorganic inputs to aquatic systems and have seen mixed results (Conley *et al*., 2009; Elser *et al*., 2007; Paerl *et al*., 2016), suggesting nitrogen is acquired from other sources. Nitrogen may be derived internally from regeneration via organic matter degradation in the water column and sediments or through nitrogen fixation. In Lake Erie, internal nitrogen regeneration is important for maintenance of *Microcystis spp.* blooms and can produce a surplus of free N (Hoffman *et al* 2022). While heterotrophic release of N is important, nitrogen fixation, a microbial mediated process in which atmospheric nitrogen (N_2_) is fixed to ammonia (NH_3_), can be a significant and underappreciated source of bioavailable ammonia in lakes (Natwora & Sheik, 2021) and in cyanobacterial blooms (Beversdorf *et al*., 2013). Together this highlights that biological nitrogen cycling is highly important to the maintenance and duration of cHABs. While, internal loading of nitrogen has long been discussed (Hampel *et al*., 2019; Hoffman *et al*., 2022), the role of nitrogen fixing organisms, both cyanobacteria and non-phototrophic microorganisms, is often-overlooked and could serve as an important supply of nitrogen to cHABs (Marcarelli *et al*., 2022; Vitousek *et al*., 2002). Furthermore, as N-limitation becomes more common (Paerl *et al*., 2018), the role nitrogen fixation plays in nitrogen availability becomes crucial, especially during bloom development, sustainment, and toxicity (Gobler *et al*., 2016; Tanvir *et al*., 2021).

Nitrogen fixation is a lynchpin in the nitrogen cycle and is important in N-limited conditions. In Lake Erie, up to 85% of N_2_ fixed through nitrogen fixation leads to N assimilation in the cell or to the surrounding microbial community (Salk *et al*., 2018). Recent work has shown that across the Laurentian Great Lakes nitrogen fixation is an important process (Natwora & Sheik, 2021). The influence nitrogen fixation has on eutrophication is especially evident in freshwater bodies that hold dense diazotrophic (nitrogen fixing microorganisms) populations that contribute to the sustainably of available nitrogen (Scott & Marcarelli, 2012; Wurtsbaugh *et al*., 2019). During cyanobacterial blooms in eutrophic systems, significant increases in gene transcription encoding the nitrogenase (*nifDKH*) protein complex suggest nitrogen fixation may be important for promoting and sustaining blooms (Lu *et al*., 2019). Nevertheless, the importance of nitrogen fixation transcends many aquatic environments and is well studied across the estuary-marine continuum (Kramer *et al*., 2018; LaRoche & Breitbarth, 2005). The long-held dogma that nitrogen fixation is inhibited by excess inorganic nitrogen does not hold true for all systems, and given nitrogen acquisition pathways and metabolism thresholds, nitrogen fixation still may be in important process in N-rich waters (Gobler *et al*., 2016; Landolfi *et al*., 2015). Given the cellular N-demand and inherent N-rich toxin compounds, nitrogen fixation may be an important microbial process that is proliferating cHABs in freshwater environments. Furthermore, the role nitrogen fixation plays in both meeting cellular N demand, and in sustaining cHABs could be especially applicable in seasonally, nitrogen limited lakes.

To address the role nitrogen fixation plays in supporting cyanobacterial bloom longevity and toxicity, we tracked microbial populations and quantified nitrogen fixation rates in the Lake of the Woods (LoW). LoW is a compelling study lake for quantifying the effect of nitrogen-fixing cyanobacteria, as it has annual toxin producing cHABs and becomes N-limited prior to the peak of the blooms (Paterson *et al*., 2017). In response to LoW’s impaired water quality status due to eutrophication (Heiskary & Wilson, 2008), research and management efforts emphasized phosphorus cycling and loading, with the goal to set a total daily maximum load (Edlund *et al*., 2017; James, 2017b, 2017a; Paterson *et al*., 2017; Reavie *et al*., 2017). While these studies presented a framework to target a single nutrient, LoW is still exhibiting cHABs despite phosphorus reduction efforts (Alam *et al*., 2020; James, 2017b). However, little work has focused on nitrogen cycling in LoW. Previous work has shown that LoW is generally nitrogen-limited during the summer growing season (Reavie *et al*., 2017). However, the presence of nitrogen-fixing cyanobacteria from the genera *Aphanizomenon* and *Dolichospermum* during peak blooms (Chen *et al*., 2009; Watson & Kling, 2017) suggests that nitrogen fixation may be an important metabolic strategy to overcome nitrogen deficient growth when waters are replete with phosphorus.

In LoW blooms, *Aphanizomenon* spp., *Microcystis* spp., and *Dolichospermum* spp. have been observed with a concomitant presence of the cyanotoxins microcystin (hepatotoxin) and anatoxin (neurotoxin) (Binding *et al*., 2011; Zastepa, Watson, *et al*., 2017). Because of seasonal N-deficient growth conditions, the ability to fix nitrogen gives *Aphanizomenon* spp. and *Dolichospermum* spp. a competitive advantage over other cyanobacteria. Interestingly, despite N-limitation in the summer both microcystin and anatoxin, which require nitrogen, are still produced. This would suggest that nitrogen recycling or fixation may be supplying necessary nitrogen for synthesizing these cyanotoxins. Given the observed dominance of *Aphanizomenon* spp. in summer/fall blooms, we hypothesized that nitrogen fixation would be an important process for the maintenance and longevity of the bloom. To address our hypothesis, we quantified nitrogen fixation rates in the surface waters of LOW during the 2021 bloom season, where the lake was experiencing periods of extreme N-limitation. We also relate these rate measurements to the total microbial and cyanobacterial community, cyanotoxin presence and abundance, and chemistry of the waters in the southern basin of LoW. Our study is the first to quantify nitrogen fixation in LoW and highlights that it is an important a source of internal nitrogen supply.

## Methods

### Station location and sample collection

Sampling efforts were focused on the southern basin of Lake of the Woods (Minnesota, USA) that encompasses Muskeg Bay and Big Traverse Bay. Lake of the Woods (LoW) is a multinational waterbody, occupying Tribal and First Nation lands and the borders of the Canadian provinces of Ontario and Manitoba and the U.S. state of Minnesota. LoW is the second largest lake in Ontario and the largest lake in the United States, excluding the Laurentian Great Lakes extending over 3850 km^2^ with a watershed area of approximately 70,030 km^2^. Lake of the Woods is an important natural, economic, recreational, and drinking water resource. LoW has inherent morphological and hydrological complexity, and noticeable alterations due to climate change (DeSellas *et al*., 2009; Schupp & Macins, 1977). Minnesota waters of LoW were declared impaired in 2008 by the Minnesota Pollution Control Agencies due to exceedance of eutrophication criteria (total phosphorus, chlorophyll*-a, and* and secchi depth) (Heiskary & Wilson, 2008). LoW consists of several distinct basins that vary in bathymetry, mixing patterns, and trophic status (J. P. Anderson *et al*., 2017). Likewise, the land use surrounding the northern and southern basins are different and consequently influence the water quality found in each (Zastepa, Pick, *et al*., 2017). The southern basin is mainly large, shallow, relatively uniform in depth, well mixed, and ranges from mesotrophic to eutrophic. The southern region is mainly surrounded by agricultural land, contributing to the productivity of the basin through runoff (Rusak & Mosindy, 1997; Zastepa, Pick, *et al*., 2017). The northern portion of LoW is surrounded by boreal forest and represents a collection of numerous islands and interconnected basins that are deeper, occasionally stratified and overall less productive than the southern basin (Binding *et al*., 2011). Cyanobacterial growth conditions have increased steadily since the 1900s (DeSellas *et al*., 2009) with increased phosphorus inputs (Pla *et al*., 2005), mean air temperatures (on average 2.5°C warmer) resulting in earlier spring ice-off dates, and changes in regional precipitation patterns (DeSellas *et al*., 2009).

Three long-term LoW monitoring stations were sampled: Muskeg Bay (48.95365, – 95.18853) in the southwest, MPCA2 (48.92182, -94.72825) located on the eastern edge of Big Traverse Bay, and BT5 (49.00002, -95.00175), a centrally located site in Big Traverse Bay (SI Figure 1). Maximum depths for the sites were 8, 9.1, and 10.4 m for Muskeg, MPCA2, and BT5, respectively. The three stations were sampled four times for nitrogen fixation rates and molecular analysis from June to October 2021 (see Figure 1 for specific dates). Nutrient and toxin samples were collected once in March, and twice monthly from June to October. Water was collected from an integrated surface composite (0-2 m) and from 1 m off the respective maximum depth using a Kemmerer water sampler. Water samples were analyzed for chlorophyll-*a*, total phosphorus, total nitrogen, ammonium, nitrate and nitrite, soluble reactive phosphorus, dissolved inorganic carbon and dissolved organic carbon (Chl-*a*, TP, TN, NH_4_^+^, NO_3_^-^+NO_2_^-^, SRP, DIC, and DOC, respectively), cyanotoxins (microcystins, cylindrospermopsin, anatoxin-a, and saxitoxin), nitrogen fixation rates (N_Fix_), and microbial community analysis using 16S rRNA gene amplicon sequencing. At each sampling station and event, a YSI Exo2 multiparameter sonde (YSI, Inc., USA) was deployed to collect vertical profiles of temperature, pH, dissolved oxygen, and phycocyanin. The volumes of water used for each water quality parameter are outlined below.

### Nutrient Analysis

Water samples were collected in acid-washed HDPE opaque bottles and stored on ice at ∼4 °C. Samples were either analyzed immediately upon return to the laboratory (SRP, DIC, DOC) or frozen at -40 °C until analysis. Water quality analyses were performed either at the St. Croix Watershed Research Station or RMB Environmental Laboratories (Detroit Lakes, MN, USA) following standard methods. Briefly, TP/SRP, TN, NH_4_^+^, and NO_3_^-^+NO_2_^-^ were analyzed following Standard Methods 4500-P, 4500-N, 4500-NH_3_, 4500-NO_3_ (APHA 2012), respectively, using a Unity Scientific SmartChem 170 discrete analyzer. DOC and DIC were analyzed using a Teledyne Tekmar Torch Combustion TOC Analyzer following Standard Method 5310-C (APHA 2012). Integrated samples were analyzed for chlorophyll-*a* concentrations via fluorometry following EPA method 445.0 (Arar and Collins 1997).

### Toxin Analysis

Water samples were collected in 50 mL glass amber vials and stored on ice until transferred to a -40 °C freezer until analysis. Algal toxins microcystin, anatoxin-a, saxitoxin, and cylindrospermopsin were analyzed using commercially available Microcystin-DM ELISA (Abraxis #52015), Cylindrospermopsin-ELISA (Abraxis #522011), Saxitoxin-ELISA (Abraxis #52255B), and Anatoxin-a RBA (Abraxis #520050) kits (Abraxis LLC, Warminster, PA) coupled with an ELISA microtiter plate reader. All toxin analyses were done per the manufacturer’s protocols at the St. Croix Watershed Research Station.

### Assessing nitrogen fixation rates using the acetylene reduction assay

Potential nitrogen fixation rates were quantified using the acetylene reduction assay (ARA) method, previously described in (Natwora & Sheik, 2021). The ARA quantifies the conversion of acetylene gas to ethylene gas by the nitrogenase enzyme (Christiansen *et al*., 2000; Stewart *et al*., 1967). Briefly, two liters of water were collected in triplicate from surface and bottom waters and concentrated on to a 0.22 μm filter (MF-Millipore Membrane filter) using a portable peristaltic pump at low pressure to minimize cell breakage. Given the dense filamentous blooms on LoW, typically only 200-500 ml was able to be filtered, due to biomass in the waters. The final volume filtered was recorded and accounted for during final calculations. Following filtration, the triplicate filters and controls were transferred to 50 mL glass serum vials, immersed in 25 mL of lake water filtrate from the corresponding sample depth, sealed with a chlorobutyl rubber stopper (Wheaton), and sealed with an aluminum cap. Acetylene gas for the assay was generated prior to field sampling by combining 10 g of calcium carbide and 100 mL of deionized water in a side arm flask and collected in a Tedlar gas sampling bag (EnviroSupply & Service). Once each sample was sealed, samples were spiked with 1 mL of acetylene gas resulting in an initial acetylene headspace of 0.15 atm (Capone & Montoya, 2001). Samples were incubated at room temperature, which were reflective of water temperatures, for 24 hours, and on the bench top to mimic natural cyclic light. After 24 hours, incubations were terminated using 5 mL of trichloroacetic acid (TCA) and stored in the dark until samples were able to be processed (approximately 1 week). Bottle headspace was sampled, and gas concentrations (acetylene and ethylene) were quantified using gas chromatography (Natwora and Sheik, 2020). Calculations of nitrogen fixation rates (nmol N_2_ L^-1^ hr^-1^) were based on methods outlined in Capone and Montoya (2001), which account for the partitioning of the acetylene and ethylene gasses between the headspace and dissolved in the medium. We used a correction ratio of 4:1 (ethylene gas to N_2_) instead of a traditional 3:1, to account for the reduced H_2_ evolution (hydrolysis) associated with the reduction of acetylene (Capone, 1993; Chia *et al*., 2019; Jensen & Cox, 1983; Tang *et al*., 2020).

### Molecular sample collection, DNA extraction, and 16S rRNA gene analysis

Two liters of water were collected from an integrated surface water composite (0-2 m) and the microbial biomass was concentrated onto 0.22 μm Sterivex filters (Millipore Sigma) using a portable peristaltic pump, again 200-500 ml of water were typically filtered due to biomass. Samples were not preserved using a commercial fluid to maximize downstream nucleic acid concentrations. Instead, filters were aseptically removed and placed in 15 mL centrifuge tubes and stored on dry ice. Samples were transferred to -80 °C and stored indefinitely until the time of DNA extraction. Total nucleic acids were extracted from the Sterivix filters using the Zymo Direct-zol DNA/RNA Miniprep kit (Zymo Research). Extracted DNA was quantified using a Qubit v3.0 (Invitrogen). Samples were shipped to the University of Minnesota Genomics Center for 16S rRNA gene amplicon sequencing. The V4 region of the 16S rRNA gene was amplified using the 515F and 806R primers (Gohl *et al*., 2016; Kozich *et al*., 2013) and sequenced using the Illumina MiSeq platform using 2x300 bp paired-end sequencing. Prior to generating amplicon sequence variants (ASVs), primers were trimmed from amplicon reads using Cutadapt (Martin, 2011). Using DADA2, amplicon reads were assessed for quality, screened for PhiX viral sequences, dereplicated, forward and reverse sequences joined, full length ASVs were generated, and chimeras were identified and removed (Callahan *et al*., 2016). Taxonomy of the ASVs was assigned to the Silva 138 (Quast *et al*., 2013) and the FreshTrain reference databases with TaxAss v2.1.0 (Rohwer *et al*., 2018). Functional capabilities of the ASVs were assigned using the Tax4Fun2 R package, that uses the metabolic capabilities of known organisms to predict the functional traits of closely related sequences (Aßhauer *et al*., 2015).

To understand the variability of toxin production genes in cyanobacterial species observed in our 16S rRNA gene libraries, we used our LoW ASV taxonomy as a guide to identify potential toxin producing cyanobacterial genera and species. We downloaded publicly available genomes from the National Center for Biotechnology Information (NCBI) (Sayers *et al*., 2022) and searched for 16S rRNA gene similarity between our ASVs and the genomes with BLASTn (Altschul *et al*., 1990). The program antiSMASH was used to identify secondary metabolite production genes in non-*Microcystis* spp. genomes, which includes microcystin, anatoxin, saxitoxin, cylindrospermopsin, anabaenopeptins, and heterocyst glycolipids (Blin *et al*., 2021). For *Microcystis* spp. we used our ASV BLASTn hits to identify genomes and cross-referenced our genome ID with the genotype traits and taxonomy from Cai *et al*. (2023).

### Data analysis methods

For all data analysis and graphing, R v4.0.4 was used in parallel with R Studio (RStudio Team, 2020). R Packages vegan (Oksanen *et al*., 2013), microeco (Liu *et al*., 2021), and ggplot2 (Wickham, 2011) were used to compute relative abundances, taxonomic classification, and graphical illustration. Prior to estimates of diversity, the ASV dataset was rarefied to 5,000 ASV sequences per sample to maximize the number of samples we could analyzed. However, as a result, the June sample from station BT5 was omitted from the final analysis.

### Data availability

All 16S rRNA amplicons are publicly available through the NCBI short read archive (SRA) project PRJNA941870.

## Results and discussion

### LoW phenology

LoW is a seasonally ice-covered lake, as such the summer growing season is dependent on the timing of ice-out in the spring and ice-in in the fall. Ice-out has varied over the last 37 years (1985-2022) with median times occurring at the first of May (n = 34, min = April 4, 2012 & 2000, max = May 21, 2014, Data from the Minnesota Department of Natural Resources). Ice-in times are also variable over the last 45 years (1977-2022) and typically occur in the last week of November (n = 9, min = November 16, 1984, max = December 2, 2019). During our 2021 sampling season, LoW’s open water season was from April 23 to November 30. Surface water temperatures peaked at ∼21 °C during the July 26 sampling time and declined over the remaining growing season (Figure 1A). Due to the shallow waters of LoW, deep-water temperatures for parts of July and August were nearly the same as surface waters (SI Figure 2B). The upper water column was oxic throughout the sampling season, and only bottom waters in July briefly reached anoxic levels (SI Figure 2). We observed two periods of phytoplankton growth quantified by Chlorophyll-*a,* May 6 – August 24 and September 15 – October 4 (Figure 1B). Our data shows there was variation between each station regarding the timing of peak bloom. However, in general the southern portions of LoW experienced peak bloom sometime during the September 21 and October 4 dates based on our measures (chlorophyll-*a*) and satellite observations (e.g., cyanoTRACKER; Mishra *et al*., 2020)

**Figure 1.**
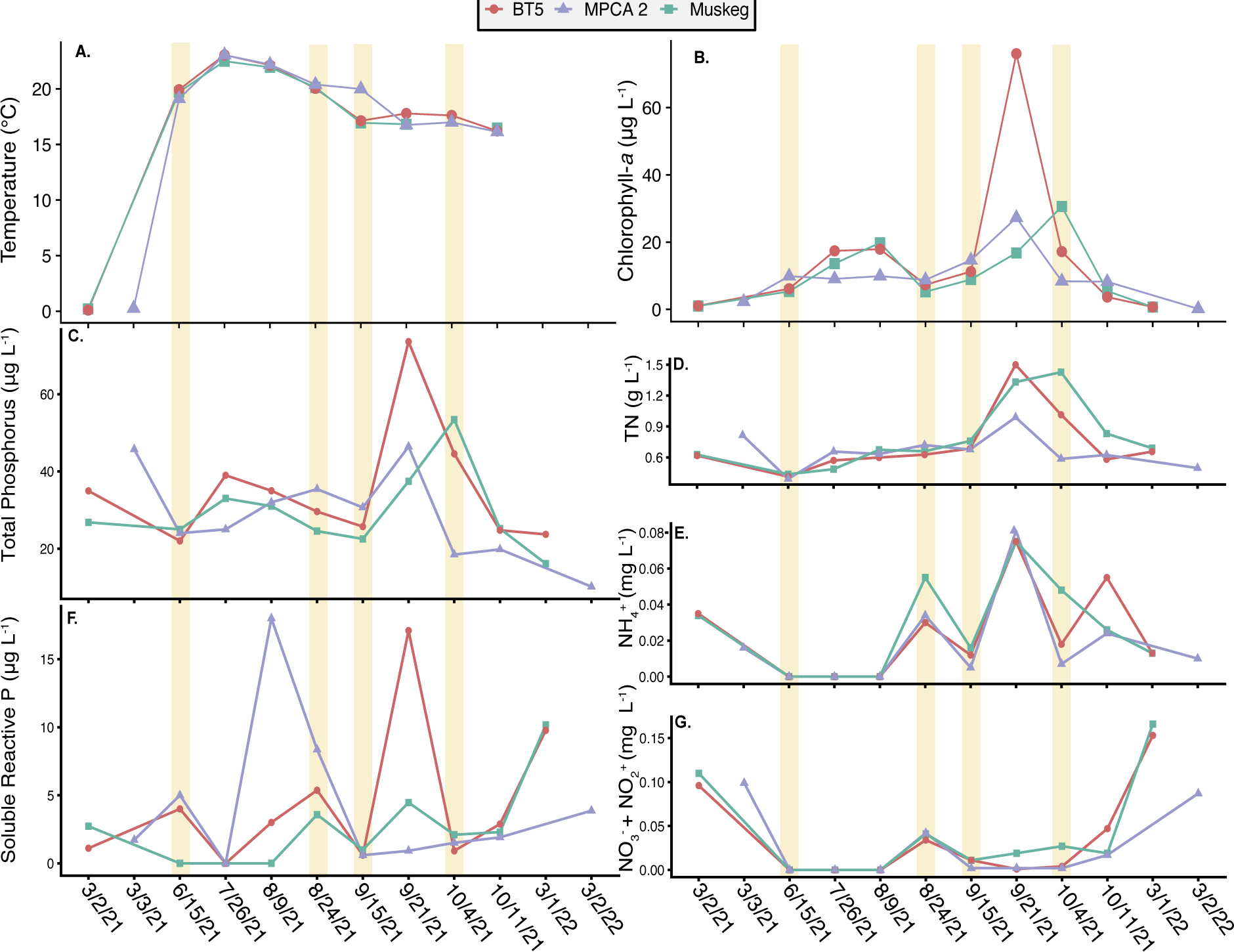
Lake of the Woods water column characteristics at the three stations during the 2021 temporal transect. A) Surface water temperatures (2m), B) Chlorophyll-*a* concentrations, C) Total Phosphorus, D) Total Nitrogen (TN), E) Ammonia, F) Soluble Reactive Phosphorus (SRP), and G) Nitrite and Nitrate (Oxidized N). Sites are represented by a red circle (BT5), purple triangle (MPCA2), and green square (Muskeg). Yellow bars represent dates where nitrogen fixation rates and DNA for microbial community analysis were collected.

### Nitrogen growth deficiency relative to phosphorus

Nutrient concentrations in LoW are dynamic and typically shift from P-deficiency to N-deficiency (Reavie *et al*., 2017). In our study we found that phosphorus, total and SRP, were detectible, much of the time above method detection limit, during each of our sampling times (Figure 1C&F). Total phosphorus was more abundant than SRP, as it contains P bound in biomass. However, the presence of SRP indicates there is a net surplus of free P throughout the LoW growing season. In contrast, during the first phytoplankton growth period (June through August), inorganic nitrogen (NH_4_^+^ and NO_2_^-^+ NO_3_^-^) concentrations were below detection limits (Figure 1E,G). Our observation is in line with previous observations (Reavie *et al*., 2017; Watson & Kling, 2017) and highlights the periods of extreme nitrogen limitations in the southern basin waters. The timing of the limitation and the increase in TN indicates that phytoplankton growth is responsible for creating and maintaining low inorganic nitrogen concentrations during this period of growth. Starting in late August, we observed several spikes in SRP, ammonia, and oxidized nitrogen concentration. The initial spike coincides with the end of the first phytoplankton growth period and suggests increases are due to either internal N cycling through biomass degradation (Gardner *et al*., 2017; Hampel *et al*., 2019; Hoffman *et al*., 2022) or a direct byproduct of biological nitrogen fixation (Xu *et al*., 2021). The second increase in chlorophyll-*a* coincides with the cyanobacterial bloom. Interestingly, oxidized nitrogen species remain low during the second growth period (Figure 1F), while Total N and ammonia increased sharply (Figure 1D). Again, the drawdown in oxidized nitrogen is likely in response to the biomass increase during the observed *Aphanizomenon* spp. bloom, as a significant increase in TN and chlorophyll-*a* is observed (Figure 1BD).

DIN to TP ratios were low throughout the peak growing season (Figure 2) and only rebound the following year during ice-in. Nitrogen-deficient growth conditions were prevalent throughout the study (N:P < 20:1, Guildford & Hecky (2000)) and are well below the Redfield ratio (<16:1, Redfield (1958)). Furthermore, during the June 15, July 26, and August 9 sampling dates, N:P ratios were likely lower than depicted, because measured inorganic N for these events was below method detection limits (Figure 2). Guildford & Hecky (2000) highlighted that in freshwater systems N:P ratios <20:1 indicate nitrogen deficient growth, ratios between 20:1 and 50:1 represent balanced growth, and ratios >50:1 indicate phosphorous deficient growth. More recently, the ratio of dissolved inorganic N to total phosphorus (DIN:TP) has been proffered as a better indicator for deducing nutrient limitation and representing *in situ* N limitation of the phytoplankton community (Bergström, 2010; Ptacnik *et al*., 2010; Reavie *et al*., 2017). Together, this indicates that phytoplankton communities in LoW are N stressed throughout the growing season and that internal cycling, either though heterotrophic processes or nitrogen fixation must be occurring to drive the observed increase in biomass during the blooms.

**Figure 2.**
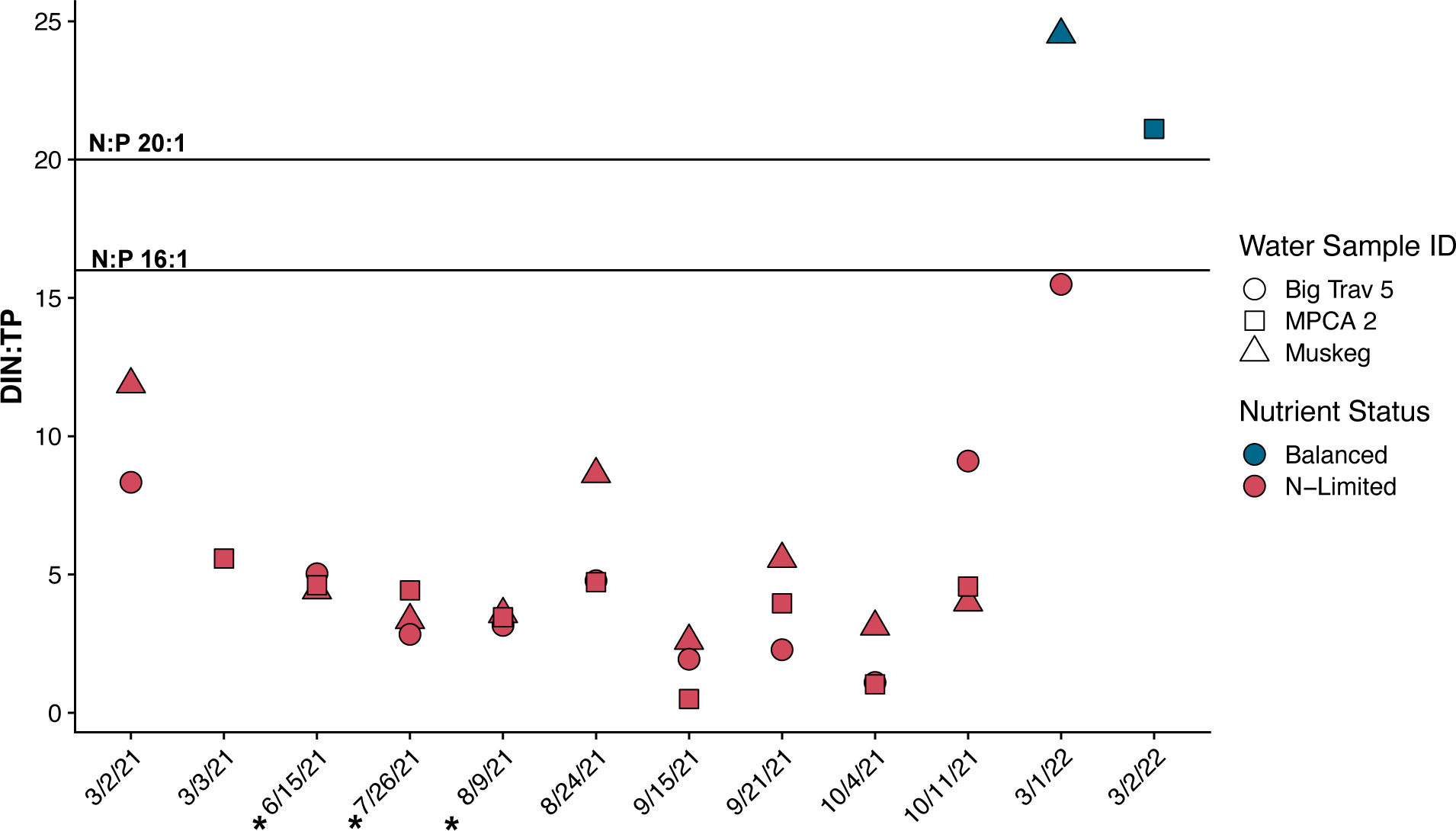
The molar ratios of Dissolved Inorganic Nitrogen to Total Phosphorus (DIN:TP) at each station during our sampling season. Shapes indicate station and fill colors represent nutrient status based on Guildford and Heckey (2004). The * indicates that inorganic N species, ammonium, nitrate, and nitrite were below detection limits. Here we have used the detection limit of the instrument to calculate a theoretical DIN:TP for these sampling dates. Horizontal lines represent Redfield (16:1) and Guildford and Hecky (20:1) nitrogen to phosphorus ratios related to growth.

### Nitrogen fixation rates increase as blooms intensify

Throughout the growing season and at all stations, nitrogen fixation rates were quantifiable and exponentially increased throughout the growing season, as the *Aphanizomenon spp.* bloom intensified (Figure 3A). In general, nitrogen fixation at all three stations were similar early in the season as the first phytoplankton growth period was beginning. Unfortunately, we were unable to sample for nitrogen fixation at every nutrient sampling timepoint. Nonetheless, as LoW transitioned to the *Aphanizomenon* spp. bloom (Mid to Late August), nitrogen fixation rates increased rapidly. Nitrogen fixation rates varied by station during the bloom and are likely due to bloom biomass differences at each station (see chlorophyll-*a* and TN concentrations in Figure 2a). Mean surface nitrogen fixation rates across all sites for June, August, September, and October across all sites were 0.109, 0.509, 1.48, and 3.72 nmol N_2_ hr^-1^ L^-1^ with standard deviations of 0.021, 0.217, 0.506, and 1.11 N_2_ hr^-1^ L^-1^, respectively. The MPCA2 station, that is furthest offshore, had the highest observed nitrogen fixation rate, 6.07 nmol N_2_ hr^-1^ L^-1^, during the October sampling despite having lower chlorophyll-*a* and TN concentrations than the other two stations. MPCA2 had very low oxidized and reduced N concentrations at the time of sampling that may have driven the high rates, as nitrogen was in high demand. Interestingly, we observed three-dates where ammonia concentrations spike (8/24, 9/21, and 10/11). The August 24 sampling appears to be during a transitional state as chlorophyll-*a* was lower than the previous samples, this would suggest that the phytoplankton community may be undergoing a turnover event and heterotrophic release of N and P are occurring. However, as the *Aphanizomenon* spp. bloom begins to intensify we see an exponential increase in nitrogen fixation rates and the depletion of oxidized N and a second spike in ammonia. This spike is likely due in part to nitrogen fixation by *Aphanizomenon* spp. The final ammonia spike is again likely due to community turnover as the bloom was waning and to nitrogen fixation rates, as they were still very high during the October-5 sampling time. This suggests that as fixation rates increase a cellular threshold is likely met and nitrogen is excreted from the cell. This physiological threshold is not well understood for most species. However, these excretion dynamics drive legume-diazotroph symbiosis (Phillips, 1980).

**Figure 3.**
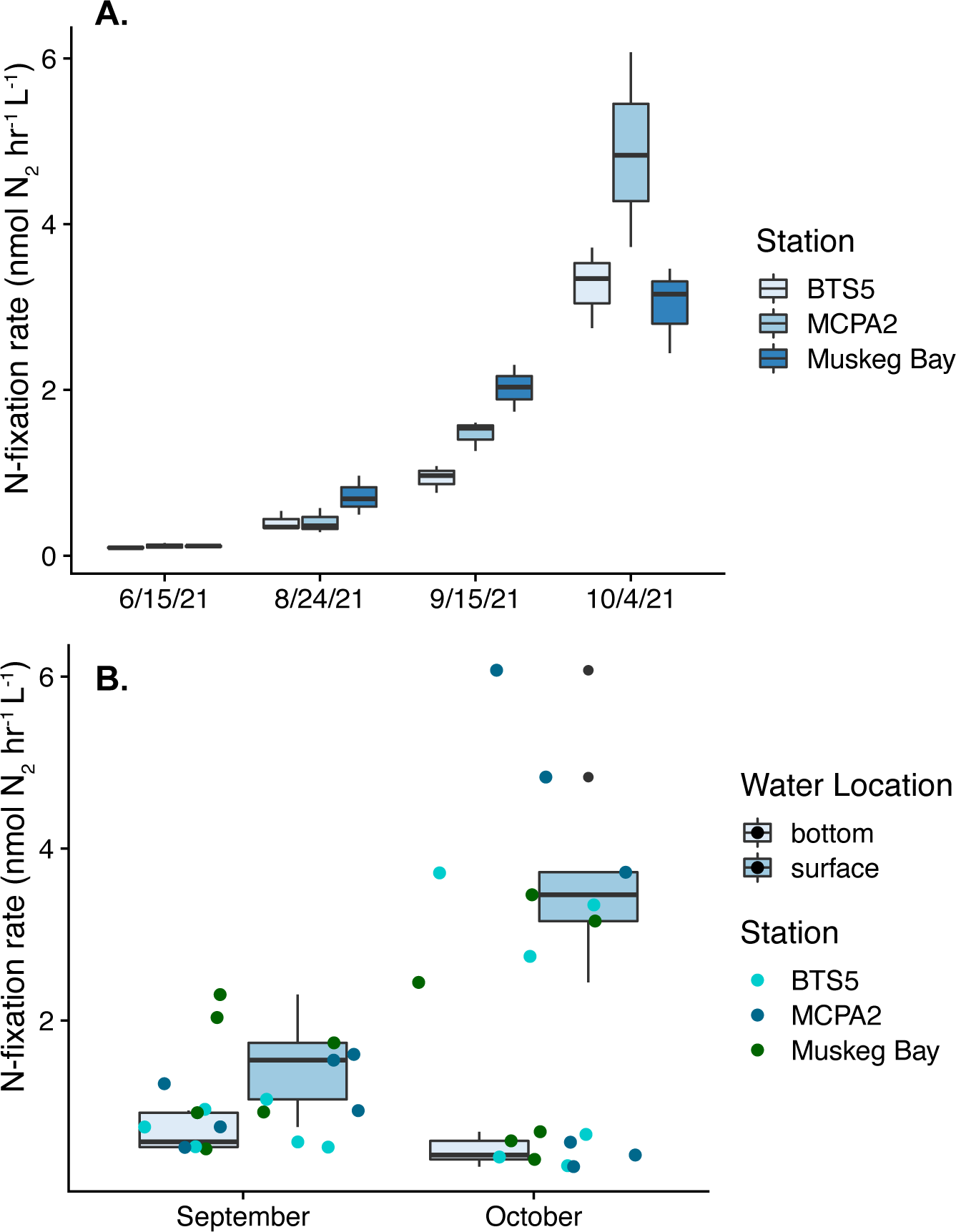
Nitrogen fixation rates at the three sampling stations (A) over the course of the 2021 summer growing season. B) Comparison of surface and bottom water nitrogen fixation rates taken in September and October during the peak of the *Aphanizomenon* spp. bloom.

As LoW is quite shallow, the potential for deep water nitrogen fixation fueling nitrogen deficiencies in the epilimnion is possible. To this end we sampled near bottom waters and quantified nitrogen fixation rates at each station during the September and October sampling times. (Figure 3B). We found that bottom water nitrogen fixation rates for September and October (mean nitrogen fixation rates 0.695 and 0.489 nmol N_2_ hr^-1^ L^-1^, respectively) were significantly lower than surface rates (p-value = 0.002), especially in October, but were still higher than surface rates from the June and August sampling (Figure 3B). We see similar depth dependent fixation rates in larger lakes (Natwora and Sheik, 2020). However, unlike deeper lakes, given the depth of the waters at our LoW sampling stations, nitrogen fixed in deep water nitrogen could make its way to the surface, especially when the waters are polymictic and well mixed (SI Figure 2). Together, internal cycling of N in LoW is complicated as it appears that both heterotrophic recycling and nitrogen fixation are operating simultaneously, though the magnitude of contribution from each process may differ temporally as the phytoplankton’s community structure changes.

### LoW cHABs primarily produce microcystin

Throughout the sampling survey, microcystin was the primary cyanotoxin identified (Figure 4). From August to October, the total microcystin concentrations were above the method detection limit of 0.15 µg L^-1^ in both surface and bottom waters for most of the sampling sites. Of the twelve sampling events, nine had total microcystin concentrations above the Minnesota Department of Health fresh drinking water advisory of 0.1 µg L^-1^, however, the recreational advisories outlined by both the EPA and Environment Canada of 8 µg L^-1^ were not exceeded. Detection of microcystins across the southern basin of LoW have been previously reported (Zastepa *et al*., 2022; Zastepa, Pick, *et al*., 2017), with a recent study reconstructing microcystin distribution and concentrations using sediment DNA dating back to ∼175 years (Pilon *et al*., 2019). We observed a peak in microcystin concentrations at all three stations during the August 24 sampling time which straddles the first and second growth periods. During this time, we also observed increases in oxidized and reduced nitrogen. We did see hints at later times that blooms at specific stations could periodically be more toxic and tied to nitrogen concentrations (September 21 and October 4). These findings are in line with Zastepa (2017) where a positive, significant correlation was observed between microcystin and nitrate (NO_3_^-^) concentrations, but not with other nutrients such as phosphorus. Together, this would suggest that nitrogen availability has a strong influence on toxin production in LoW.

Less abundant cyanotoxins were detected during our sampling including cylindrospermopsin, anatoxin and saxitoxin (SI Table 1). Cylindrospermopsin was observed near the detection limit (0.05 μg/L) at every station and nearly all timepoints. We also saw variable presence of anatoxin and saxitoxin in our samples, however, both were below reportable detection limits (0.15 μg/L and 0.02 μg/L) and could represent non-specific substrate targeting from the Abraxis assays. A recent study also reported detection of anatoxin-a and cylindrospermopsin in surface samples collected on LoW. The detection of these toxins cooccurred with detection of the necessary genes responsible for synthesis, *ana* and *cyrA*. Although saxitoxin was not detected in their study, the *sxtA* gene responsible for saxitoxin production was detected through qPCR. Detection of *sxtA* occurred at lower frequencies than an important microcystin gene, *mycE,* representing ∼78% respectively, but still more than half, ∼59% respectively (Zastepa 2023).

**Figure 4.**
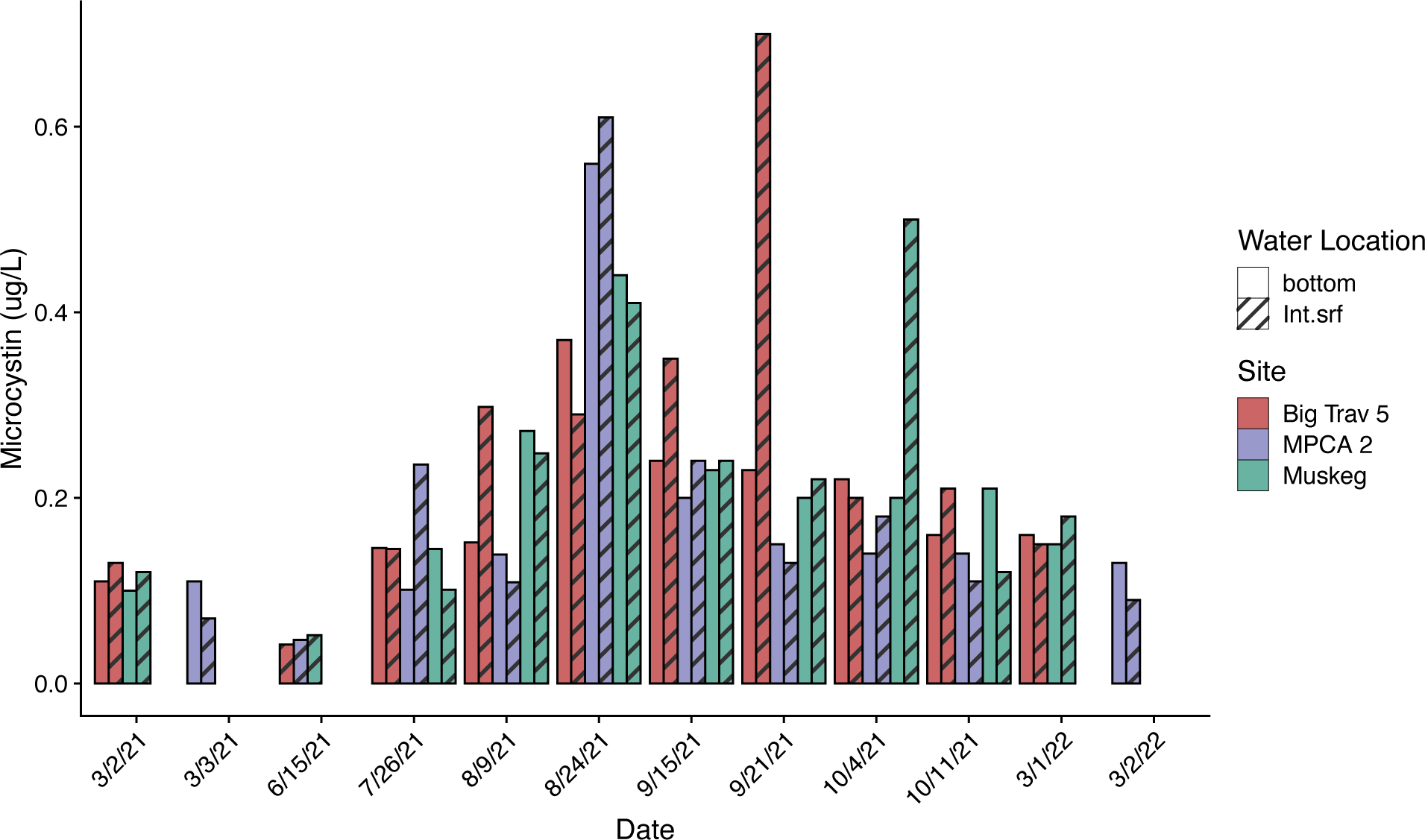
Microcystin detection at each station (depicted by color) over time. Integrated surface waters (Int.srf) are represented with a hatched bar.

### 16S rRNA gene amplicons reveal dominance of Aphanizomenon during the bloom

Throughout the 2021 sampling season, non-photosynthetic and potentially heterotrophic bacterial phyla were more abundant than Cyanobacteria (Figure 5A), which is consistent with other blooms (Berg *et al*., 2009; Buchan *et al*., 2014; Sheik *et al*., 2022). The abundance of heterotrophic microorganisms highlights their multifaceted roles as they are stimulated by the productivity in blooms but their presence and growth represents an important sink for phosphate and ammonium (Kirchman, 1994), however, they are essential for the recycling of nutrients like ammonia (Hampel *et al*., 2019). At all-time points, the bacterial community composition (BCC) was comprised primarily of the phyla Cyanobacteria, Bacterodia, Gammaproteobacteria, Verrucomicrobiae, Alphaproteobacteria, Actinobacteria, and Planctomycetes (ranked in descending order of abundance; Figure 5A). In June, Bacteroidia, Gammaproteobacteria, and Alphaproteobacteria comprised much of the microbial community but decreased relative to Cyanobacteria as the summer intensified. Broadly speaking, Phyla, such as Bacteroidia, Proteobacteria, Actinobacteria, and Verrucomicrobia, all contain organisms that can specialize in macromolecule degradation (i.e., complex carbon, proteins/amino acids, simple sugars, etc.). Thus, their importance and that of the heterotrophic community (Archaea, Bacteria, and micro/macro-Eukaryotes) cannot be understated (Pound *et al*., 2021; Smith *et al*., 2022). Together, the heterotrophic community helped drive the observed spikes in nitrogen and phosphorus observed over the course of the study that contributed to the spur in cyanobacterial bloom growth.

**Figure 5.**
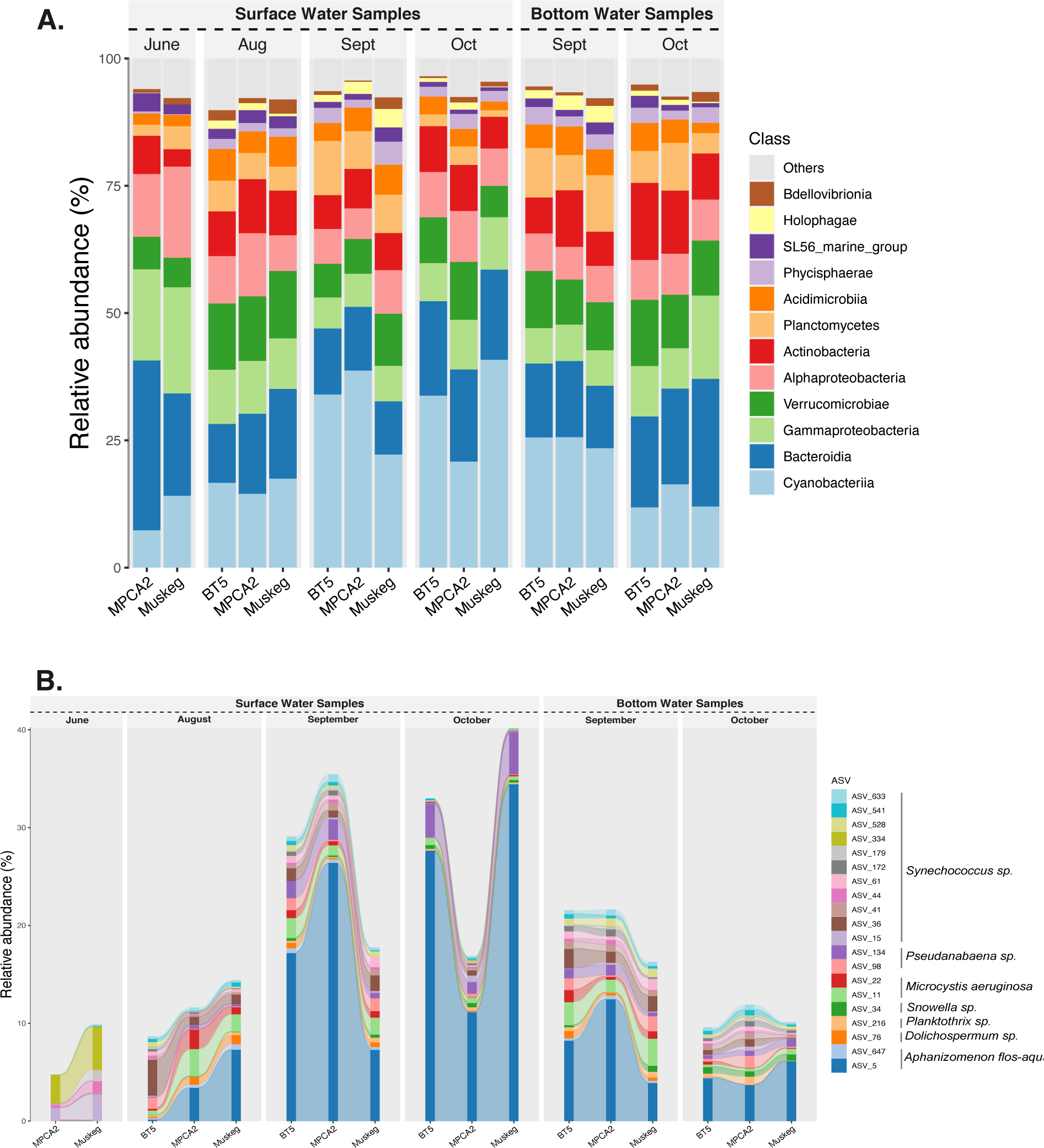
Bacterial community composition of the A) whole community at the Class taxonomic rank and B) the top twenty Cyanobacteria amplicon sequence variants (ASV) abundance during our study. Taxonomy follows the Silva 16S rRNA gene database version 138.

Cyanobacteria were less abundant in June and primarily comprised of four *Synechococcus* ASVs (Figure 5B). Upon reaching near uniform water column temperatures (July – August, SI Figure 2), cyanobacteria become the dominant taxa observed, comprising roughly 15% of the total microbial community. The shift in abundance is commiserate with typical cyanobacterial growth, as water temperatures and light availability increase (Havens *et al*., 2003; Rohwer *et al*., 2023). The diversity of dominant cyanobacterial ASVs also increase as we see *Dolichospermum, Snowella, Microcystis, Aphanizomenon,* and different *Synechococcus* ASVs all being present and at different abundance based on sampling station (Figure 5B). By our September sampling, nitrogen-fixing *Aphanizomenon* spp. (Figure 5B), as previously observed (Watson & Kling, 2017), were the dominant cyanobacteria present and represented by a single ASV. Other less abundant nitrogen-fixing cyanobacterial genera were also present, including *Dolichospermum* (Liang *et al*., 2020), *Pseudanabaena* (Pasteur Culture Collection of Cyanobacteria, 2000*), Nodosilinea* (Perkerson III *et al*., 2011) and *Synechocystis* (Brass *et al*., 1994). Using Tax4Fun, the most prevalent nitrogen fixing bacteria were Cyanobacteria, however, we also identified nitrogen fixing ASVs associated with Alpha- and Gamma-proteobacteria and the Gemmatimonadales (SI Figure 3).

Within the top 20 ASVs, we also saw non-nitrogen fixing cyanobacterial genera, *Planktothrix*, *Microcystis*, and *Snowella.* Although previously thought that P ultimately limited phytoplankton biomass in LoW (Watson & Kling, 2017), the physiology and emergent properties of the shifting community suggest nitrogen plays a crucial role in structuring the microbial community. Similarly, eutrophic lakes, Mendota and Utah, transition from a non-diazotrophic to a diazotrophic dominated microbial community in response to dissolved inorganic nitrogen availability (Beversdorf *et al*., 2013, H. Li *et al*., 2020). In the case of Lake Utah, cyanobacterial communities shift from picocyanobacterial *Cyanobium* to filamentous *Aphanizomenon* and *Dolichospermum* (H. Li *et al*., 2020). Moreover, a significant increase in nitrogen fixation gene expression was observed pre-bloom vs bloom conditions (H. Li *et al*., 2020). Together, these suggest that nitrogen concentration, availability, and the diazotrophic community are important for promoting and sustaining cHABs.

We observed striking exponential increases in nitrogen-fixation rates as the bloom intensified. The dominance of nitrogen fixing *Aphanizomenon* were likely the main drivers of this spike in rates. While nitrogen fixation can supply bioavailable nitrogen to the cyanobacterial cells, this process is leaky, as fixation rates likely exceed growth requirements. Thus, the surrounding community may benefit from the nitrogen fixing organisms and influence their growth and abundance (Vitousek & Howarth, 1991). The possibility that the diazotrophic community could support non-nitrogen fixing cyanobacteria like *Microcystis* and *Planktothrix* was explored in Lake Utah. The study suggested that excess N, fixed by diazotrophic cyanobacteria like *Aphanizeomon* and *Dolichospermum*, could also be supplying the nutrients promoting the growth of *Microcystis* and *Planktothrix* (H. Li *et al*., 2020). On a cellular level, N is essential for cell maintenance and regulation of cellular homeostasis (Forchhammer & Selim, 2020). Many studies have explored the ideal stoichiometric ratio of C:N:P needed for cellular maintenance, and the molar ratio that delineates nutrient deficiencies in the environment (Bergström, 2010; Guildford & Hecky, 2000; Ptacnik *et al*., 2010; Redfield, 1958). However, some cyanobacteria are stoichiometrically plastic depending on the availability of nutrients (Ji *et al*., 2020; Osburn *et al*., 2021). Together, this highlights that as LoW experiences N-limitation, nitrogen fixing organisms increase abundance concomitantly as fixation rates increase. Thus, supplying the nitrogen needed to meet cellular demand of themselves and potentially other organisms within the community.

### Potential toxigenic cyanobacterial species in LoW blooms

At LoW, we identified, at low concentrations, a suite of cyanotoxins– microcystin, saxitoxin, anatoxin, and cylindrospermopsin. While microcystin is commonly observed in cyanobacterial blooms, the presence of saxitoxin, anatoxin, and cylindrospermopsin are less common and understudied, and may be increasing in frequency in freshwater systems in North America (Christensen & Khan, 2020; Cruz *et al*., 2013; Testai *et al*., 2016). With the continuous production of cyanotoxins at LoW, the likelihood of long-term, low dosage, exposure may have broad health impacts on those who rely on LoW for drinking water (Yang *et al*., 2022; Zhao *et al*., 2020). Thus, from a management perspective it is important to understand who is responsible for each toxin produced to develop mitigation strategies.

Our 16S rRNA gene survey identified several candidate cyanobacterial genera that may produce cyanotoxins. Using our ASV sequences, we used BLASTn to identify putative reference cyanobacterial genomes. All LoW ASVs were highly similar (>98%) to reference genomes and in some cases multiple ASV sequences were associated with a single genome (Table 1). The presence of multiple ASVs associated with the same genome likely originates from genomes containing multiple, slightly different 16S rRNA gene sequences (Pei *et al*., 2010). Using antiSMASH (Blin *et al*., 2021), we identified whether these genomes contained genes for cyanotoxin production and thus could potentially produce cyanotoxins (Table 1). We found five genomes associated with microcystin and three genomes associated with either anatoxin, saxitoxin, or cylindrospermopsin. The genomes of the microcystin candidates consist of two *Microcystis spp*., two *Dolichospermum spp.*, and one *Planktothrix rubescens*. The two *Microcystis* genomes belong to the newly identified taxonomic clusters P (M049S1) and 19 (M048S1) (Cai *et al*. 2023). All genomes in these two *Microcystis* clades contained the genes for microcystin production. The two *Dolichospermum* genomes both fall within a single GTDB species (*Dolichospermum* sp000312705, also known as *Dolichospermum lemmermanii*), but associate with different cultured strains, UHCC 0352 and OL01. The presence of microcystin genes within this GTDB species is quite variable (Driscoll *et al*. 2017, Osterholm *et al*. 2020, Sheik *et al*. 2022), and they have previously only been found in European strains (Sheik *et al*. 2022). The presence of microcystin production genes in the OL01 strain, which was isolated from Odell Lake, Oregon, USA (Dreher *et al*., 2021), is the first genome from this species isolated in North America to potentially produce microcystin and suggests it may be more widespread than previously thought. This genome is also very similar to the genome recovered from the 2018 bloom in Lake Superior that was not found to carry microcystin production genes (Sheik *et al*., 2022). From our analysis it is unclear if the LoW *Dolichospermum* sp. is capable of microcystin production and either a cultivation, metagenomic, or metatranscriptomic approach is needed to confirm. Finally, we also identified a *Planktothrix rubescens* genome that contains genes for microcystin production. *Planktothrix* spp. are common bloom formers across the Laurentian Great Lakes, especially in Lake Erie’s Sandusky Bay (Davis *et al*., 2015; Rinta-Kanto & Wilhelm, 2006; Salk *et al*., 2018) and microcystin production is common across this genus (Pérez-Carrascal *et al*., 2019). *Planktothrix* are efficient nitrogen scavengers (Hampel *et al*., 2019; McKindles *et al*., 2022) and some can fix nitrogen (Pancrace *et al*., 2017).This would suggest that they would be effective bloom forming species at LoW. However, these five genomes represent a small percentage of the total cyanobacterial population at most times sampled, except for *Microcystis. Microcystis* ASV’s 21 and 11, were quite abundant during the August sampling (Figure 5b) when microcystin concentrations and free nitrogen also were quite high (Figure 4 and Figure 1). However, we should note that *Planktothrix spp.,* can produce more microcystin per cell than *Microcystis spp*. (Fastner *et al*., 1999). Thus, may still be a significant source of microcystin despite their low abundance.

As previously suggested (Watson & Kling, 2017; Zastepa, Watson, *et al*., 2017) the LoW *Aphanizomenon* spp. are likely not capable of producing microcystin. We found an absence of microcystin genes in all publicly available *Aphanizomenon flos-aquae* genomes (SI Table 2. GTDB taxonomy *Dolichospermum flos-aquae*). However, the *Aphanizomenon flos-aquae* genome we identified was potentially capable of producing saxitoxin (Table 1). Saxitoxin, was detected but below reporting thresholds throughout the course of our monitoring. The high abundance of *Aphanizomenon flos-aquae* (ASV_5) during the bloom would suggest that if they can produce saxitoxin, concentrations should be much higher. The presence but not abundance of saxitoxin, suggests the primary *Aphanizomenon flos-aquae* strain we observe may not be capable of saxitoxin production or that saxitoxin production is highly regulated. It is understood that Lake Erie blooms are cohabitated by toxin-producing and non-toxin-producing *Microcystis aeruginosa* (Akins *et al*., 2020; Lei *et al*., 2015). Thus, similar dynamics may also occur in other cyanobacterial genera and species. The dearth of free nitrogen at most samplings, likely plays an important role for regulating the production of all nitrogen-rich cyanotoxins in LoW. Saxitoxin compared to microcystin has a very low C:N ratio (1.4 vs 4.9, respectively). At LoW, the combination of high nitrogen demand of these compounds and the observed nitrogen growth deficiencies (Figure 3) likely constrains the cell’s ability to produce these nitrogen rich compounds in high concentrations. When free nitrogen was measured in non-growth periods (Figure 1, August and October), we saw increases in microcystin (Figure 3) and saxitoxin (SI Table 1). Thus, from a management perspective, nitrogen inputs to LoW need to be limited, as fertilization will likely increase toxin production.

We identified both anatoxin and cylindrospermopsin periodically throughout our sampling. Like saxitoxin, they were below reporting thresholds. We identified two candidate genomes that could produce anatoxin or cylindrospermopsin. We found ASV_947 was identical to *Dolichospermum* sp001277295 (*Anabaena sp.* 54), which antiSMASH predicted to contain anatoxin production genes. This species/strain was isolated from Lake Saaskjarvi, Finland but there are also several genomes and cultures that have been isolated from North America, and potentially carry anatoxin production genes (SI Table 2). *Aphanizomenon flos-aquae* has been reported to produce cylindrospermopsin and has been implicated as the likely cyanobacterium responsible for cylindrospermopsin production in LoW (Preussel *et al*., 2006). However, we identified only ASV_1242 as being a potential candidate for cylindrospermopsin production.

This ASV was identical to *Dolichospermum* sp017355425 (*Dolichospermum* sp. DET66). These genomes within this GTDB species all derive from cultures collected from Detroit Lake, Oregon, USA. Together, these results highlight that waters of LoW contain a suite of cyanobacteria, both abundant and rare, that may produce cyanotoxins. The nitrogen demand of these cyanotoxins likely help constrain the production in LoW as N-limitation is pervasive. However, it should be recognized that the presence of emergent, low abundance cyanobacteria may become problematic under the right suite of environmental conditions.

ASV sequence similarity to publicly available genomes. Presence of toxin genes were identified using AntiSMASH (non-*Microcystis* spp.) or from Cai *et al*. (2023).

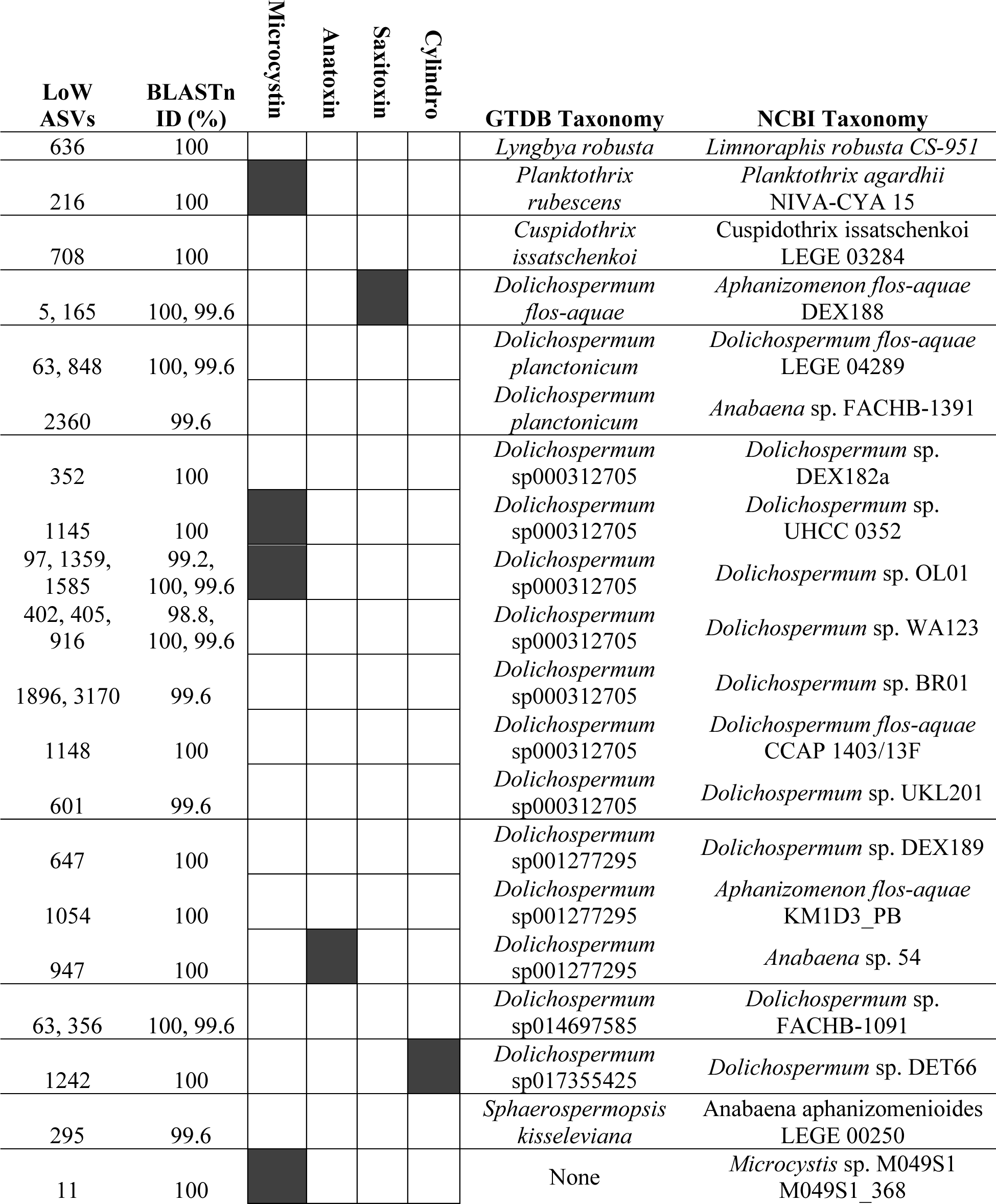
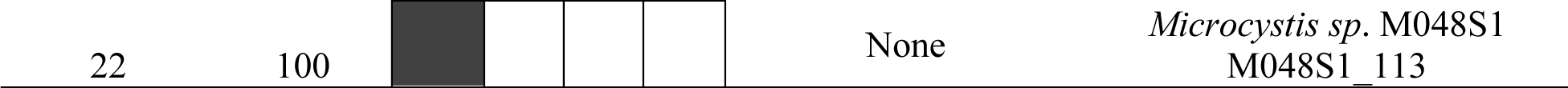

### Implications of nitrogen availability aiding the development and toxicity of LoW blooms

The dual role of nitrogen, in both controlling growth and toxicity has been explored (Gobler *et al*., 2016; Tanvir *et al*., 2021). However, research on the role nitrogen fixation plays in fueling cyanotoxins is limited. Many studies show a positive correlation with cyanotoxin production and nitrogen availability. Specifically, *Microcystis aeruginosa,* both in single-strain laboratory experiments and in the field, responds to increased nitrate concentrations with a concomitant increase in microcystin toxin quota (Gobler *et al*., 2016; Horst *et al*., 2014). Likewise, ammonium and dissolved organic N also have been linked to higher microcystin toxin concentrations in freshwater lakes, supporting the importance nitrogen has on toxin production (Monchamp *et al*., 2014). However, at LoW, microcystin is produced at relatively low concentrations despite limited inorganic nitrogen being present. This suggests that nitrogen fixation, internal recycling of nitrogen (Gardner *et al*., 2017; Hampel *et al*., 2019) or likely both may supply nitrogen to produce these nitrogen-rich toxins. Lake Mendota, like LoW, supports reoccurring *Aphanizomeonon flos-aquae* blooms in the late summer to fall months when the basin becomes nitrogen limited. Furthermore in Lake Mendota, *Microcystis aeruginosa*, is suspected to be responsible for microcystin production, as the *Aphanizomeonon flos-aquae* likely is not capable of microcystin production (Beversdorf *et al*., 2013; Zastepa, Watson, *et al*., 2017). It was suggested that the N inputs from the diazotrophic community were large enough to support *Microcystis aeruginosa* growth and microcystin production (Beversdorf *et al*., 2013). Thus, given the nitrogen state and cyanobacterial community structure in LoW, it is likely a similar dynamic is taking place, and the dominate nitrogen fixing *Aphanizomenon* spp. could support non-nitrogen fixing cyanobacterial growth and toxin production. Like Lake Mendota, microcystin production is likely associated with *Microcystis* spp., however, we also saw several cyanobacterial species that could produce it. We observed that *Microcystis* was a small proportion of the total cyanobacterial population at nearly all sampling timepoints, apart from the August 24 samples (Figure 5B), where we saw a concomitant spike in abundance, microcystin, and nitrogen concentrations. This would suggest the depleted nitrogen pool in LoW limits total microcystin concentrations by limiting the growth of microcystin producing cyanobacteria, and from a management perspective, nitrogen additions to LoW could have a damaging effect on the basin’s toxicity.

## Conclusions

Our work suggests nitrogen availability plays a large role in structuring the freshwater microbiome, importantly the cyanobacterial species composition, and further interdependently influences nutrient dynamics and secondary metabolites. Our results show that during periods of nitrogen limitation, the microbial community is mainly composed of the nitrogen-fixing genus *Aphanizomenon*, whose metabolic activity is likely supporting in part the less abundant microcystin producing cyanobacteria. In addition, the use of molecular techniques reveals that most cyanotoxins concentrated in LoW are likely produced from low abundance toxigenic cyanobacteria. For most of the season, LoW was nitrogen deficient, and at times, inorganic N could not be quantified; however, microcystins, a nitrogen rich compound, were still being measured. Together, this suggests the bioavailable nitrogen produced from nitrogen fixation could be used to help support the microcystin producing cyanobacterial community, and potentially shuttled into toxin production. Given that it is likely the nitrogen environment in LoW is structuring the microbial community composition and influencing toxin production, further monitoring and control of inorganic N will be important for management implications. Ergo, by changing the nitrogen environment, the N:P stoichiometry could ultimately change the cyanobacterial composition in LoW, thereby altering the relative abundances of toxigenic cyanobacteria. As revealed in our ASVs, many toxigenic cyanobacteria are in low abundances, compared to *Aphanizomenon,* and are likely responsible for the cyanotoxins detected in Lake of the Woods. Altering the N:P ratio in LoW could result in many of those toxigenic species that are at low abundance to bloom or increase metabolic activity and increase the cyanotoxin production in LoW.

## Acknowledgements

This work was funded through grants to AJH from the Red Lake Nation and the Minnesota Pollution Control Agency. The molecular analysis was internally funded through the Sheik Lab and did not receive any specific grant from funding agencies in the public, commercial, or not-for-profit sectors. We thank the Polaris Foundation and Polaris Marine for providing and supporting the R/V *Navicula*. We thank the SCWRS staff Alaina Fedie, Jeremy Howland, Heather Rankin, Erin Mortenson, Amelia Wilson-Jackson for their lab and field support.

## Author Contributions

KEN, AJH, MBE and SEB collected field samples necessary for measuring all parameters outlined in this manuscript. KEN, AJH, MBE, SEB analyzed all water sample parameters outlined in this manuscript. JDC provided molecular analysis and support generating graphics and figures. KEN and CSS drafted the manuscript. AJH and MBE helped finalized and edit the manuscript. KEN, AJH, MBE, JDC and SEB approved the final manuscript.

