## Supplemental Information for "Nitrogen dynamics and fixation control cyanobacterial abundance, diversity, and toxicity in Lake of the Woods (USA, Canada)"

+1-218-726-8128

orcid: 0000-0003-0413-1924

### SUPPLEMENTAL FIGURES

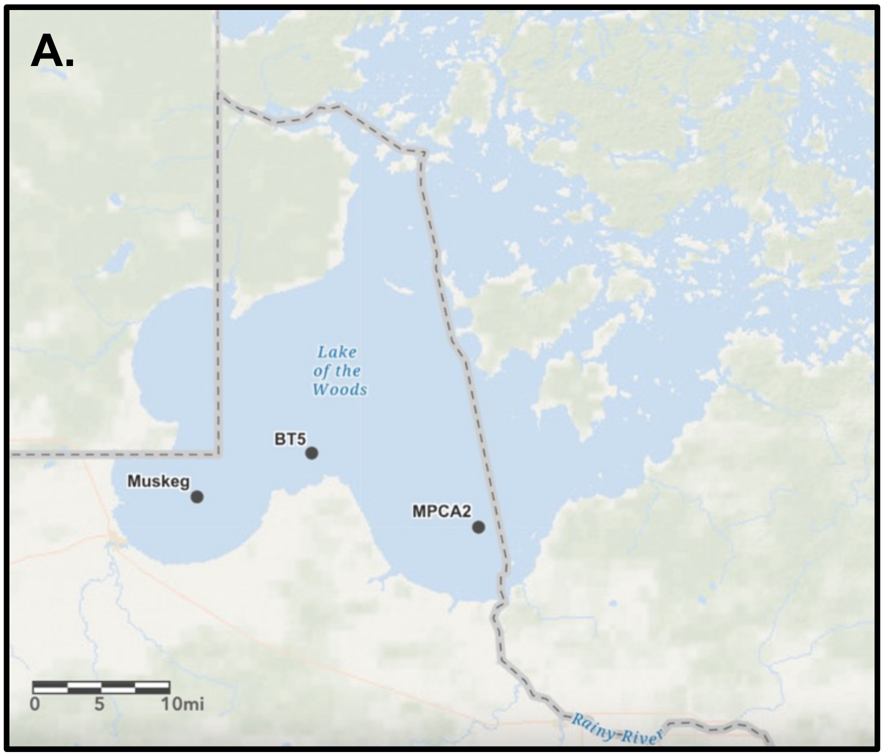


**SI Figure 1**. Locations of long-term sampling stations within Lake of the Woods.

### SUPPLEMENTAL FIGURES

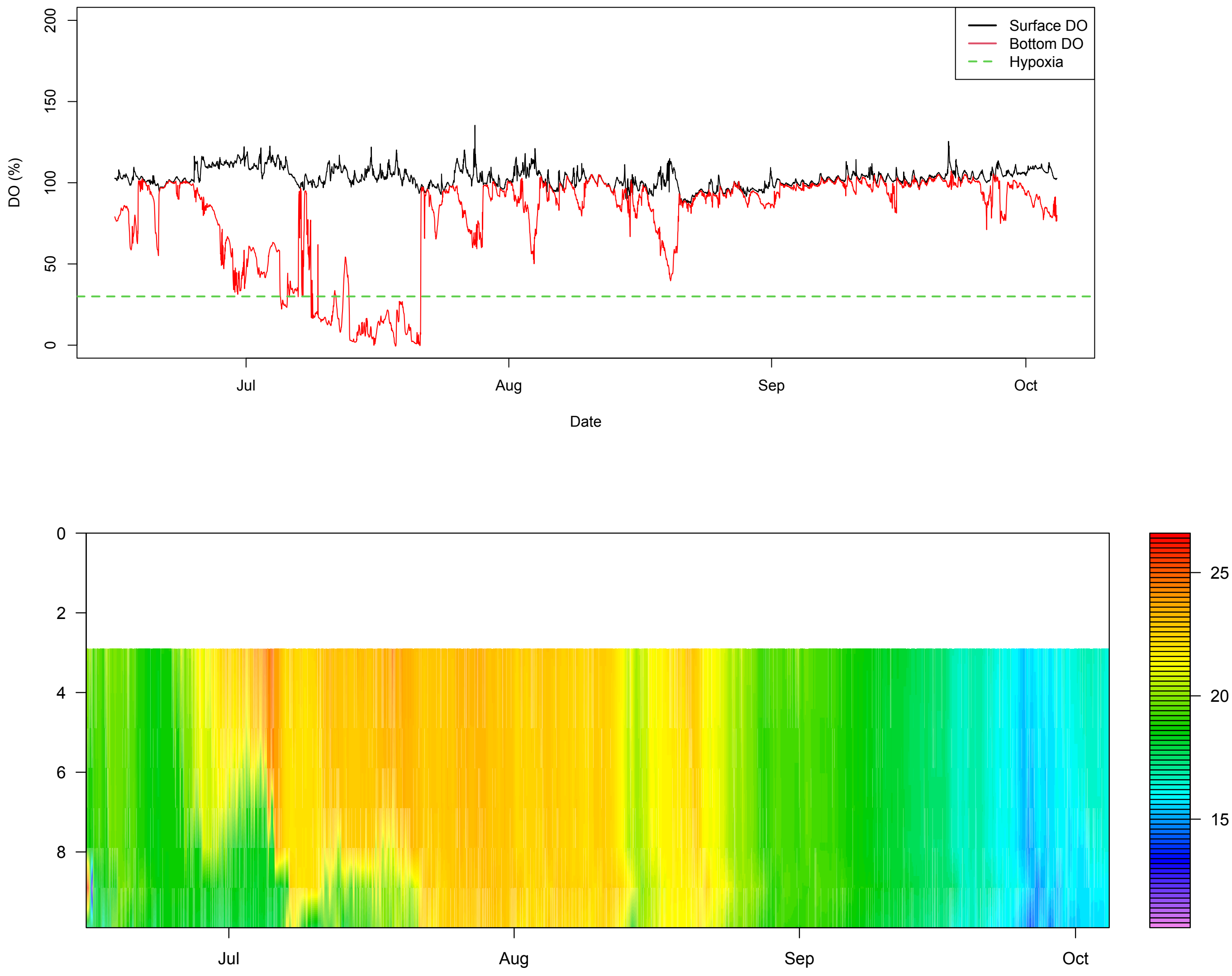


**SI Figure 2.** Dissolved oxygen concentrations (upper panel) and integrated temperature profiles from the BT5 station.

SUPPLEMENTAL FIGURES

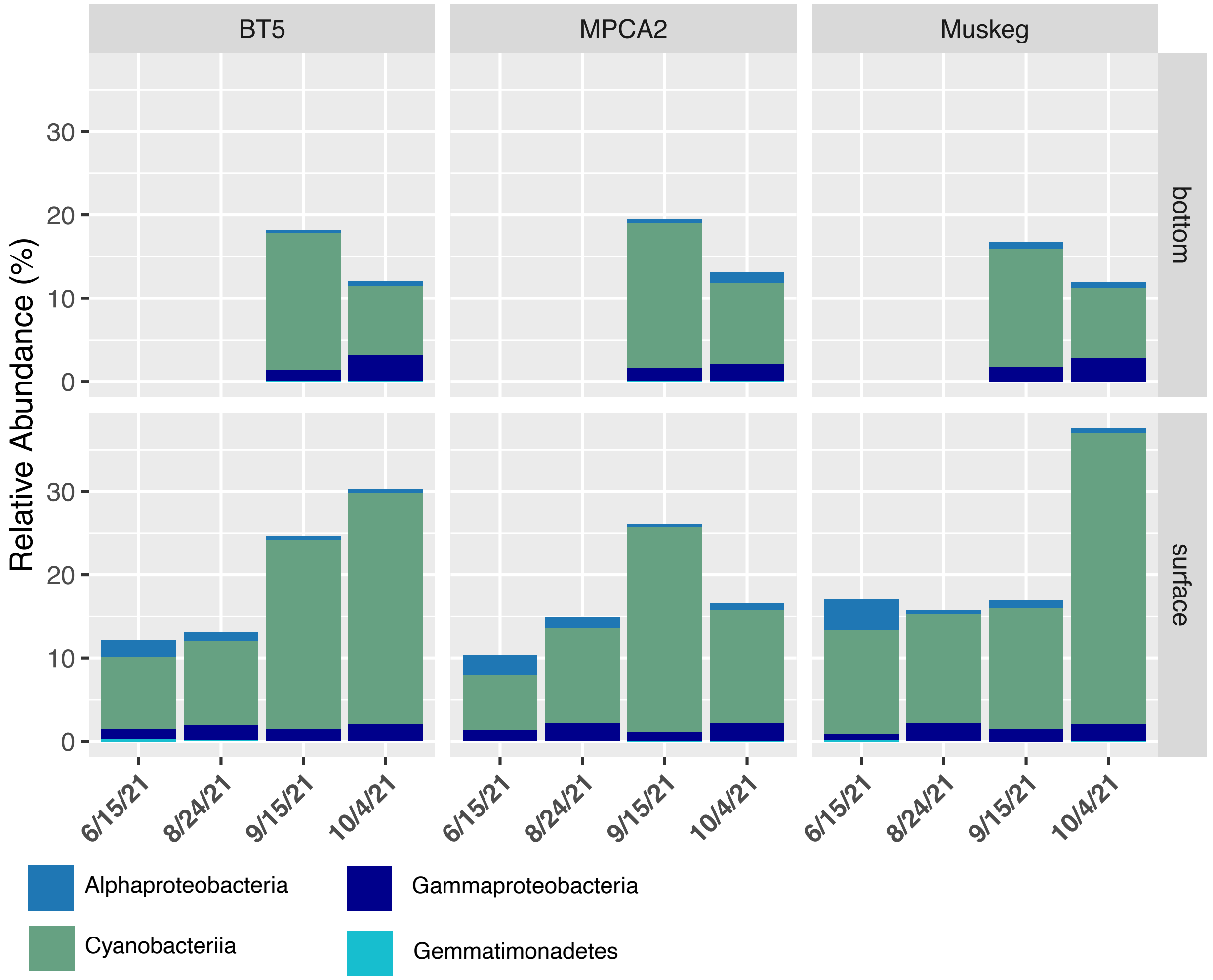


**SI Figure 3.** The predicted abundance of nitrogen fixing microorganisms using Tax4Fun, which uses custom 16s rRNA gene databases to identify sequences closely related to microorganisms with known metabolic function.
